# Running reduces firing but improves coding in rodent higher-order visual cortex

**DOI:** 10.1101/214007

**Authors:** Amelia J. Christensen, Jonathan W. Pillow

## Abstract

Running profoundly alters stimulus-response properties in mouse primary visual cortex (V1), but its effects in higher-order visual cortex remain unknown. Here we systematically investigated how locomotion modulates visual responses across six visual areas and three cortical layers using a massive dataset from the Allen Brain Institute. Although running has been shown to increase firing in V1, we found that it suppressed firing in higher-order visual areas. Despite this reduction in gain, visual responses during running could be decoded more accurately than visual responses during stationary periods. We show that this effect was not attributable to changes in noise correlations, and propose that it instead arises from increased reliability of single neuron responses during running.

---

To understand perception, it is important to study how contextual variables affect the representation of sensory information in neural populations. Locomotion, a highly ethological behavior in rodents, has been shown to have pronounced effects on the magnitude and consistency of responses to visual stimuli^1–9^. However, the investigation of these effects has so far been limited to primary visual cortex (V1). Recent work has shown that in V1, firing rates increase^2^, response variability decreases^4,9^, noise correlations decrease^8^, and signal-to-noise ratio (SNR) increases^8^ during bouts of running. These observations lead to the prevalent view that running acts to enhance visual representations both in firing rate and coding accuracy^1–4,8,10^. A normative theory that accounts for this enhancement proposes that running triggers a visual selective-attention mechanism, as vision is the most navigationally-important sense for rodents^11^. An alternate theory posits that the goal of visual cortex is to predict the next frame of visual stimulus (as in predictive coding^12^), a framework in which access to the speed at which an animal is navigating the environment is essential. Knowledge of the impact of running speed modulation on higher order, functionally specified visual regions is unknown, and might either support or reject these hypotheses.

Using the Allen Institute Brain Observatory^13^ dataset, we quantified how running speed affects visual responses in six visual cortical regions: primary visual cortex (V1), lateral visual cortex (VISl) (`LM', or secondary visual cortex), posterior medial visual cortex (VISpm) (a putative ventral stream region^14^), anterior lateral visual cortex (VISal), and anterior medial visual cortex (VISam), rostral lateral visual cortex (VISrl) (putative dorsal stream regions^14^), in cortical layers 2/3, 4, and 5 (Fig 1a). This dataset allowed us to assess the tuning of hundreds to thousands of cells in each region and cortical layer (Supplementary Fig 1a, b). A substantial fraction of neurons in all regions and layers were tuned to running speed (Fig 1e, f), and the total percent of neurons tuned to running in each region was highly predictive of the average correlation coefficient between running and neural data in that region (Figure 1i, j). The distribution of running speed modulation differed across cortical layers; layer 5 had the largest fraction of neurons tuned to running, while layer 4 had the smallest (Fig 1h). The relative paucity of neurons tuned to running in the input layer and relative abundance tuned to running in the output layer supports the view that running speed modulation is not inherited from thalamic inputs, and may originate in cortex. This is consistent with reports that neurons in LGN are not strongly modulated by running speed^2^.

**Figure 1.**
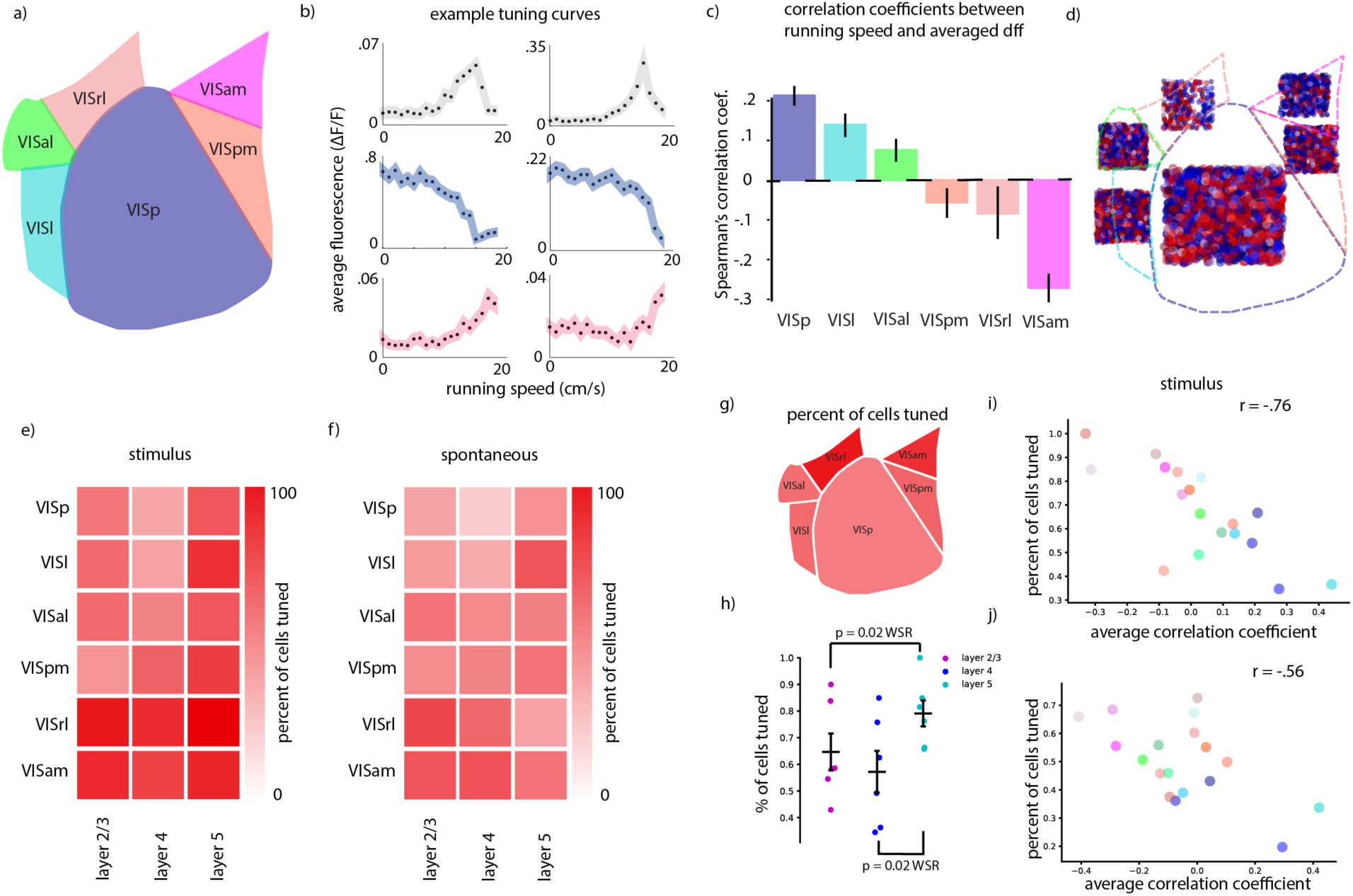
Schematic of regions included in study. **b**. Example non-monotonic, monotonically decreasing, and monotonically increasing tuning curves to running speed, top to bottom. **c**. Spearman's correlation coefficient between fluorescence (df/f) and mouse running speed. Error bars are SEM over individual imaging experiments. Data from layer 2/3, 4, and 5 are combined in each region. **d**. Spatial distribution of neurons of different tuning types, with blue (red) representing neurons with decreasing (increasing) running-speed tuning. Squares not to scale between regions. **e**. Overall fraction of neurons significantly tuned to running in each region and Cre-line, calculated during visual stimulus presentation. **f**. Same as e. except calculated in the absence of visual stimuli. **g**. Visualization of spatial distribution of overall tuning to running in the visual regions. **h**. Comparison of percent of neurons tuned in layer 2/3, layer 4 and layer 5 (significance computed by Wilcoxon signed rank test). Each data point is a different visual region. **i**. Correlation between percent of cells tuned to running and Spearman's correlation coefficient between running and average df/f. Each data point is grouped over all mice in an individual region and cre-line. **j**. Same as i. except calculated in the absence of visual stimulus.

In all areas we examined, we found a diversity of tuning to running speed, including neurons with monotonically increasing, neurons with monotonically decreasing, and neurons with significant but non-monotonic tuning to running speed (fig 1b). Although a large proportion of the neurons were significantly tuned to running speed, we found that in higher order visual cortices (especially VISam, VISpm, and VISrl), running tended to be suppressive, meaning that increased running speed *decreased* neural firing rates (fig 1c). This suppressive modulation was consistent across cortical layers, and was also present in experiments where running-speed tuning was calculated when no stimulus was present (Supplemental figure 1g). These trends were also present when we separately analyzed data from natural and artificial visual stimuli (Supplementary figure 1e,f). In general, more anterior extra striate regions had both a higher fraction of cells tuned to running, and a higher fraction of cells suppressed by running (figure 1d, g). These data did not address whether this observed reduction in firing simply reflected lower baseline firing rates during arousal^8^. Nevertheless, the observed suppressive effects of running on firing rates contradicts the naïve hypothesis that running induces selective attention to vision that increases the gain of responses throughout visual cortex, or that increased gain throughout cortex is a hallmark of mechanisms for disambiguating self-motion from object motion.

Previous studies have shown that, in addition to its effects on firing rates, running increases the fidelity of visual responses as measured by the accuracy with which visual stimuli can be decoded^8,15^. This effect has been attributed to both increases in firing rates and decreases noise correlations during running^15^. We sought to test whether, despite their reduced firing rates, increased decoding performance was still present in higher order visual areas.

To assess decoding performance, we trained a linear classifier (using multinomial logistic regression) to decode which of 8 different grating orientations was presented to a mouse. Decoding performance was significantly higher for responses during running than for responses during stationary periods (figure 2a). This trend was also present in most individual visual region and Cre-lines we investigated, despite the fact that in many of these regions firing rates decreased during periods of locomotion. Indeed, when we excluded from all datasets any neurons whose firing rates were enhanced during locomotion, we observed the same enhancement of decoding performance (figure 2b).

**Figure 2.**
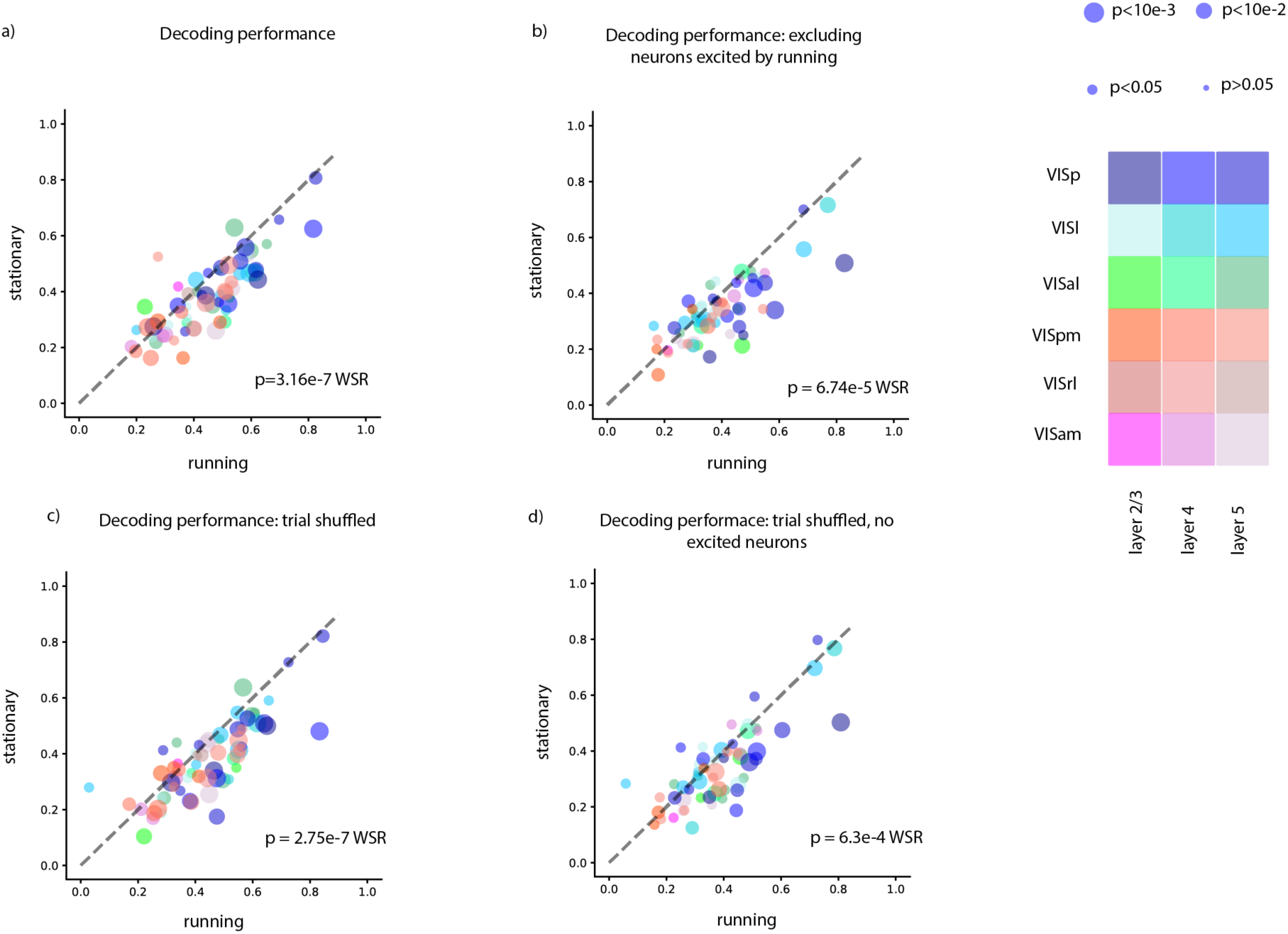
Decoding performance (multinomial logistic regression) during running and stationary periods. **a**. Average fraction of correctly classified visual stimuli during running and stationary periods (average over ten 50:50 train/test splits). Each data point is an individual experiment. Colors indicate brain region recorded; size of dot indicates significance level for difference between running and stationary decoding in an individual experiment. **b**. Same as a. but excluding neurons that increase their firing rate during running. **c**. same as a. but trial-shuffled to remove noise correlations. **d**. same as a. but excluding neurons that increase their firing rate during running and trial-shuffled to remove noise correlations. All statistics Wilcoxon signed rank test.

To investigate what changes in neural response statistics led to this improvement in classifier performance, we analyzed the noise correlations of population responses. Previous work has shown that noise correlations in V1 decrease during running^7,8^, and this has been thought to be a primary reason for improved decoding accuracy during running compared to during stationary periods^15^. Additionally, increased behavioral discrimination performance in mice during cholinergic modulation (which is typically present during locomotion) has been attributed to de-correlated neural firing patterns^3^. To determine whether decreased noise correlations during running epochs were responsible for the increases in classifier performance we observed, we compared decoder performance on data that were trial shuffled. Surprisingly, we found that — although trial shuffling slightly reduced the size of the improvement in decoding accuracy during running — a robust difference in decoder performance between running and stationary periods persisted for shuffled data (figure 2c). We were nevertheless curious whether a combination of increased response gain during running and reduced noise curious could account for the difference in decoding performance, as has been previously suggested^15^. We therefore repeated our decoding analysis on a dataset that was trial shuffled after removing all neurons excited by running; surprisingly, the difference in decoding between running and stationary periods was still present (figure 2d). These trends were consistent across multiple choices of classifier (Supplemental figure 2).

Motivated by these findings, we hypothesized that individual neurons might encode the stimulus identity more reliably when the animal was running, even if their firing rates did not increase. Indeed, we found that neurons whose firing rates did not increase during running showed increased 'reliability', defined as the variance of each neuron’s average response to each image divided by the total variance of each neuron’s response^10^ (figure 3a-c). This improved reliability was correlated with the increased decoding performance (figure 2k). Thus, unlike previous reports based on data from V1, we found increased decoding accuracy correlated with increased fidelity in single neuron responses, and was not explainable entirely by decreased noise correlations or increased stimulus responsivity.

**Figure 3.**
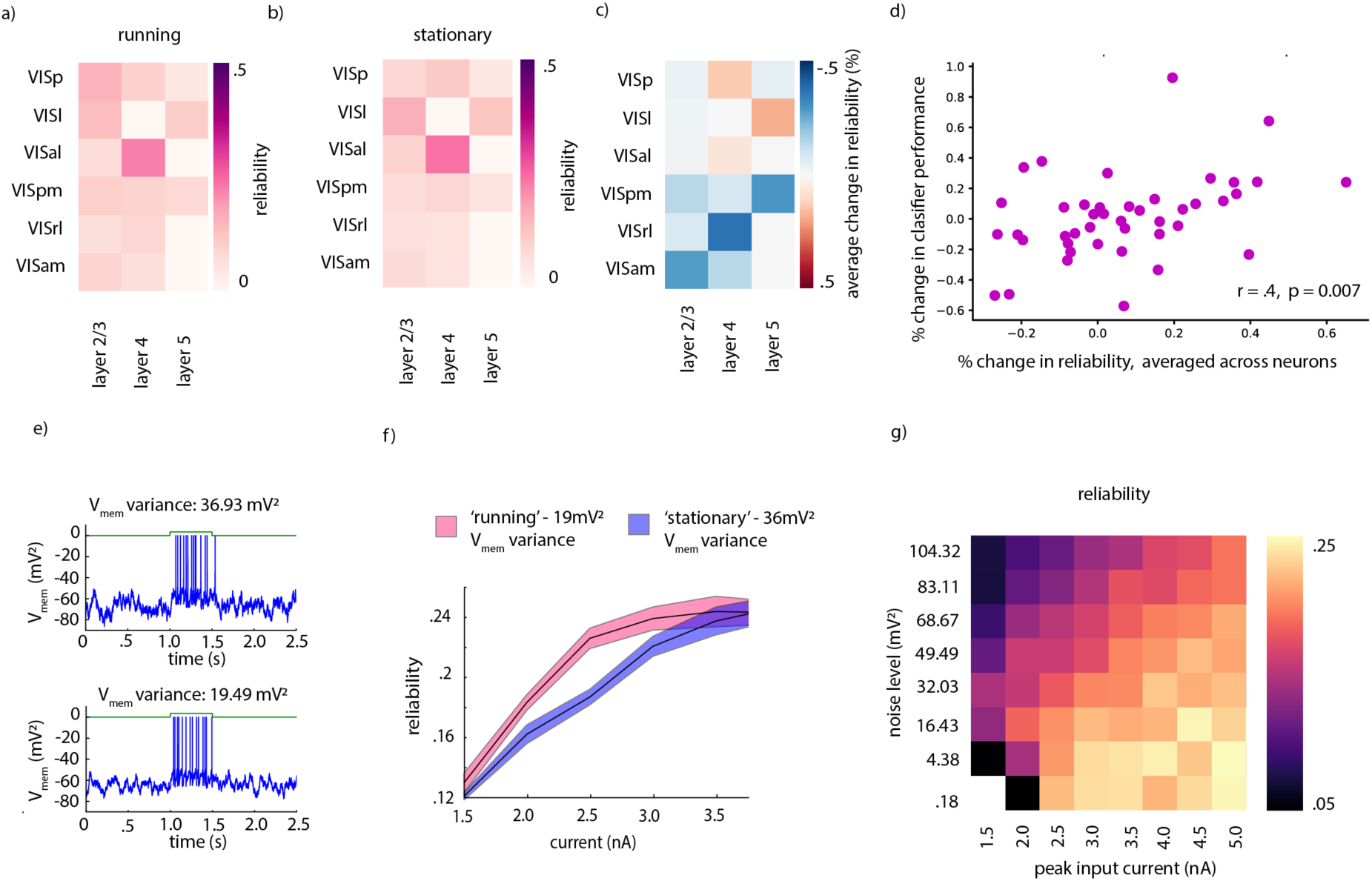
**a**. Average reliability of neurons in each brain region and Cre line when the mice are running, excluding neurons that are excited by running. **b**. Same as a. except during times when mice are stationary. **c**. Relative change in reliability between running and stationary periods. **d**. Correlation between decoding performance and average percent change in reliability in each experiment. **e**. LIF simulation with additive Gaussian noise corresponding to membrane voltage measured during stationary (top) and running (bottom) epochs **f**. Reliability versus input current for running and stationary membrane voltage noise levels. **g**. Reliability versus membrane voltage and peak input current.

Lastly, we sought to investigate possible physiological mechanisms underlying the increased reliability of single-neuron responses during running. It has been previously reported that during periods of locomotion, background membrane voltage fluctuations of neurons in V1 decrease^8,9^. We performed simulations of leaky integrate and fire neurons (LIF) (figure 3e) to determine whether this decreased membrane voltage fluctuation could counteract the expected reduction in reliability in neurons whose firing rates either decreased or did not change due to running. We added Gaussian noise to the membrane voltage of LIF neurons, while driving them with input current drawn from Gaussian shaped tuning to 15 different objects. As expected, we observed that increasing the peak input current increased the neuron's firing rate and its reliability, and that adding noise to the membrane potential also increased the neuron's firing rate, but reduced its reliability (figure 3k). We chose two noise levels reflective of levels measured *in vivo* (19 mV^2 and 36 mV^2) for running and stationary epochs respectively^9^, and simulated responses across a range of peak input current amplitudes. We found a sharp increase in reliability of these simulated responses between noise variances of 36 mV^2 and 19 mV ^2, implying that neurons had significant room to decrease their firing rates while still improving reliability. In combination with the observation that lowered background noise itself can lead to lower firing rates without changes the mean synaptic drive to a neuron, our simulations explain how neurons firing rates could easily be reduced by ~50% during locomotion, while response reliability nonetheless increased. Further experimentation and physiological measurements will be required to establish whether a reduction in membrane voltage fluctuations during locomotion explains the enhancement in decoding performance we observed, however our simulations are consistent with this hypothesis.

In conclusion, we observed a striking difference in the type of firing rate changes during locomotion across different visual cortical regions. Surprisingly, neurons in all layers of higher order visual areas were more likely to be suppressed than enhanced by running, in contrast to previous results showing response enhancement V1. This suppressive tuning is not easily reconcilable with theories that explain the running speed modulation in V1 simply by enhanced 'attention' to vision during running, or by the need to disambiguate self-motion from the independent motion of visual stimuli. However, despite the trend towards running-induced suppression in higher order visual cortices, running still enhanced the representations of visual stimuli, as measured by decoding accuracy. This effect was not attributable to noise correlations or enhanced sparsity, but instead was attributable to increased reliability of individual neural responses. Physiologically, this could result from lower background membrane voltage fluctuations during locomotion. These results highlight previously unknown differences in running speed modulation in primary visual cortex and higher order visual cortices, and should inform future theories that seek to explain the function of the observed modulation of visual signals during locomotion.

## Supplementary Information

**Supplemental Figure 1.**
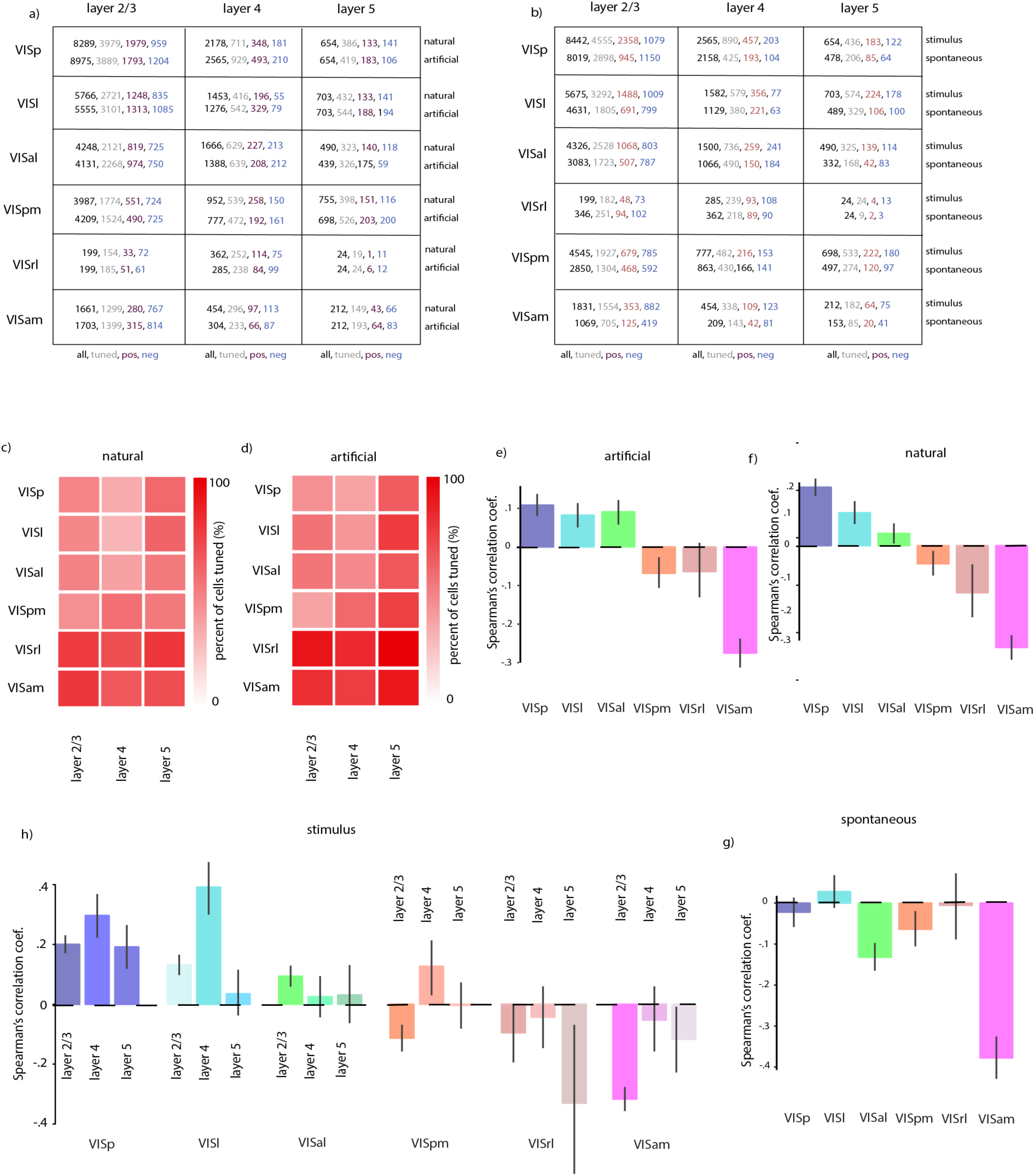
Number of tuned neurons displaying monotonic increasing vs. monotonic decreasing tuning to running, split out by natural and artificial stimulus types. b. Number of neurons tuned to running, split out by periods with and without stimulus. c. Percent of neurons tuned to running, natural stimuli. d. Percent of neurons tuned to running, artificial stimuli. e. Pearson's correlation coefficient between running speed and dF/F in each region, calculated only artificial stimuli (e.g. gratings, noise) were displayed. f. Same as e. but calculated only when natural stimuli (natural scenes, natural movies) were displayed. g. same as e. but calculated only when no stimuli were presented. h. Same as e. but with individual cre-lines displayed, calculated across all stimuli types.

**Supplemental Figure 2.**
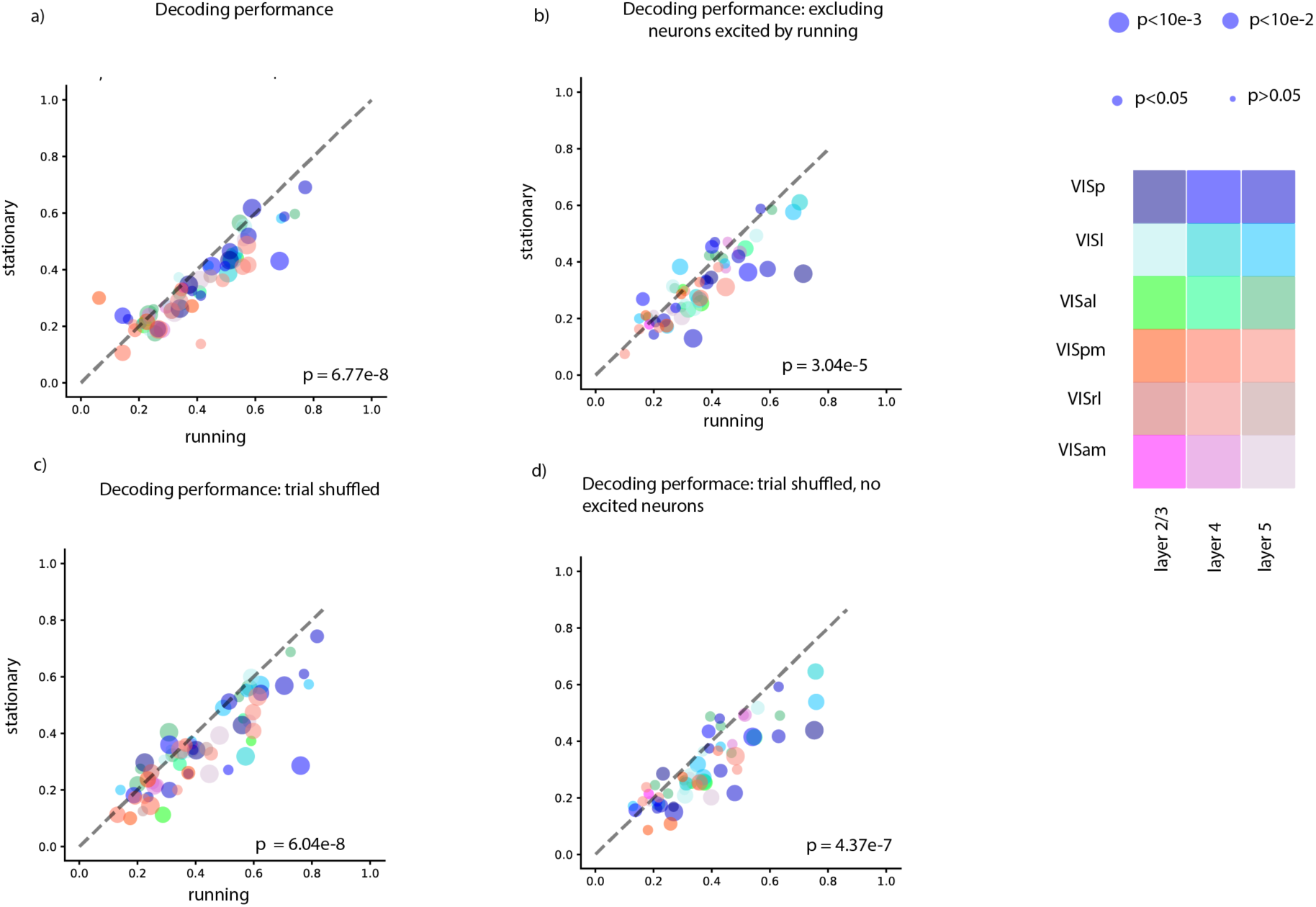
Average fraction of correctly classified stimuli across 10 CV splits, during running and stationary periods. Each data point is an individual recording experiment, colors indicate from which brain region data were recorded, size of dot indicates whether the difference between running and stationary decoding performance was statistically significant in an individual experiment. b. Same as a. but excluding neurons that increase their firing rate during running. c. same as a. but trial shuffling to remove noise correlations. d. same as a. but excluding neurons that increase their firing rate during running and trial shuffling to remove noise correlations. All statistics Wilcoxon signed rank test. Decoder: Gaussian Naïve Bayes.

**Supplemental Figure 2a.**
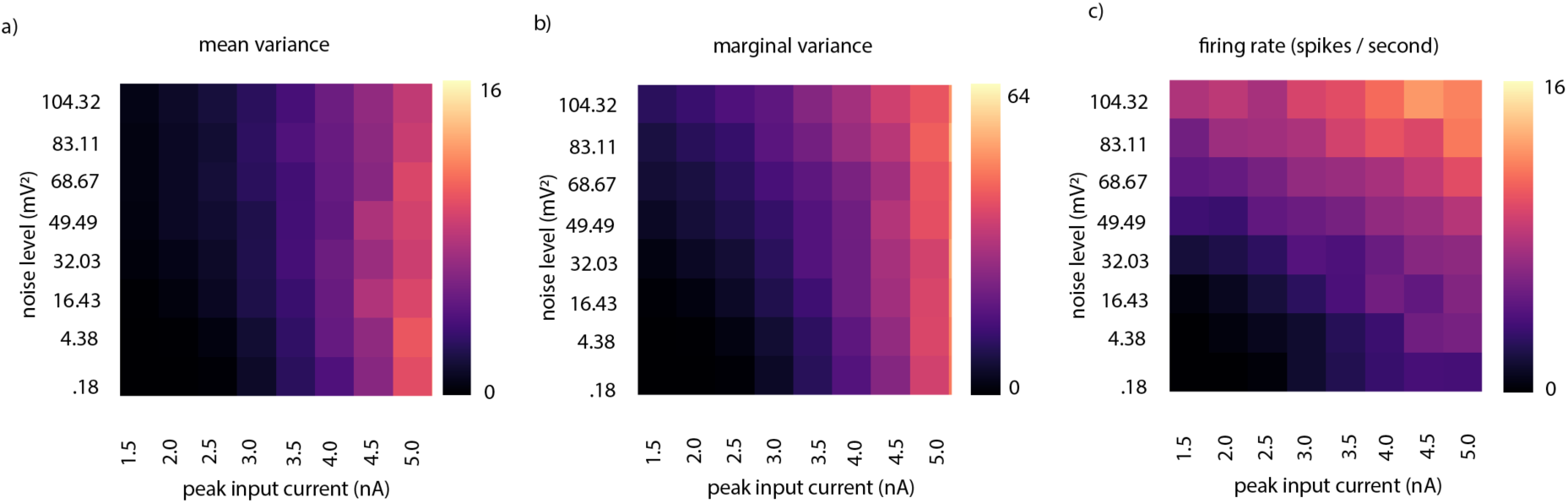
Variance of class means versus noise level and peak input current in LIF simulations. b. Marginal variance versus peak input current and noise level in LIF simulations. c. Firing rate versus noise level and peak input current in LIF simulations.

## Methods

### Data Collection

We analyzed data from the publicly available Allen Institute for Brain Science Brain Observatory data set. Their full data collection methodology can be found in the white paper^13^. In brief, transgenic mice expressing GCaMP6f in laminar-specific subsets of cortical pyramidal neurons underwent intrinsic signal imaging to map their visual cortical regions before cranial windows were implanted above the desired visual region. Mice were habituated to head fixation before imaging sessions in which they were shown either natural scenes, natural movies, locally sparse noise, or gratings. Neuropil corrected fluorescence change (df/f) traces for each cell were extracted using automated, structural ROI based-methods (See Allen Institute white paper for details). We performed no pre-processing on df/f traces after downloading through the AllenSDK. Eye movements and locomotion speed were recorded; locomotion speed is available as part of the publicly available data set. We analyzed data from the Cux2-Cre-ERT2 (layer 2/3), rbp4-Cre (layer 5), and Rorb-Ires-Cre (layer 4) mice, as data from these transgenic lines were available for all regions in the data set.

### Running speed tuning

For our analyses of running speed tuning, we selected mice who ran for at least one quarter of the stimulus presentation period (to ensure enough data-points to accurately calculate tuning). We also excluded mice whose maximum running speed was less than 15cm per second, to ensure enough of a range of running speeds were present to accurately assess correlations. We estimated running-speed tuning curves by first binning data into 20 equal-sized bins (i.e., 20 quantiles) of running speed, ranging from zero to the maximum speed attained by each mouse, and taking the mean neural activity in each bin, effectively creating a non-parametric ‘tuning curve’ to running. To determine whether a neuron’s firing was significantly modulated by running, we compared the neuron’s running-speed tuning curve to a running-speed tuning curve computed from randomly permuted data using Levene's t-test of variance; we considered a neuron tuned if its non-shuffled tuning curve had significantly more variance than it's shuffled tuning curve^5^. We calculated Spearman's rho on the binned data, to determine whether each neuron was monotonically tuned to running. We considered neuron with a rho < 0 to be suppressed by running, and a rho > 0 to be enhanced by running. We used an alpha-level of p = 0.05 for our estimate of significant tuning. We calculated this tuning both separately for natural (natural scenes and natural movies), artificial (drifting gratings, static gratings, and noise stimuli), and spontaneous activity. We found that tuning was similar across natural and artificial stimulus conditions, and therefore grouped them together in the main text (but see Supplemental figure 1).

### Decoding analysis

We performed all decoding analyses on the Drifting Gratings dataset from the Allen Institute Brain Observatory. Details of this dataset can be found in their white paper^13^, but briefly: each mouse was presented with 75 repetitions of 8 drifting gratings of different directions, for 2s per presentation, with a 1 second blank period in between stimuli. Each grating presentation had a spatial frequency of 0.04 cpd and a temporal frequency randomly selected from a set of 5 different temporal frequencies. We performed decoding of grating direction while ignoring temporal frequency. For the purposes of the decoding analysis, we excluded individual experiments in which fewer than 10 neurons were recorded – this exclusion criteria mainly applied to lower levels of VISrl in the analyses when considering all neurons except those with positive tuning to running. Note that decoding analyses included data from additional mice that were excluded from the speed tuning curve analyses (due to insufficient time spent running). Decoding analyses that excluded neurons with positive tuning to running, however, only included data from the subset of mice whose tuning curves had been well characterized.

To perform decoding, we extracted a vector of neural population activity for each trial by averaging fluorescence (df/f) over a 2s window that was offset by 330 ms (10 imaging frames) from the beginning of stimulus presentation. This time window was chosen by selecting the window that maximized the R^2^ prediction performance of held out trials from the PSTH. To compare decoding during running vs. during stationary periods, we split the data into “running trials” (trials with average velocity > 3cm/s) and “stationary trials” (trials with average speed < .5 cm/s). We randomly sub-sampled the data to ensure equal number of trials per visual stimulus class in both running and non-running subsets. For the data presented in the main paper, we performed decoding of neural responses using an 8-way multinomial logistic regression (MLR) classifier, as implemented in the scikit-learn^16^ python package. Classifier weights were learned via the LBFGS algorithm, and no regularization was applied (in separate cross-validation experiments we determined that neither L1 or L2 regularization significantly improved classifier performance, data not shown). Classifier performance was assessed via a cross-validation procedure: fraction of correctly labelled stimuli on a test set comprising 50% of the data was averaged over 10 random (class balanced) train-test splits. For shuffling analyses, we randomly permuted each neuron’s responses across trials (within the same class) so that population response vectors contained non-simultaneous responses, breaking trial to trial correlations between simultaneously recorded neurons. In the figures, decoders were both trained and tested on shuffled data, although in separate analysis we either only tested or only trained on the shuffled data, without significantly different results. The main results are consistent across different decoder types, with results obtained with Gaussian Naïve Bayes presented in the supplemental materials.

We assessed each cell’s reliability, defined as the variance of the average df/f response across different stimuli divided by the total variance across responses to all stimuli^10^. Thus, if a cell has high variance of response across all stimuli (e.g. a greater dynamic range in its responses to different stimuli types), with low trial to trial noise, it is considered to be extremely reliable – thus, a single measurement of that neuron’s response contains a large amount of information about stimulus identity.

### Leaky Integrate and Fire Simulations

To examine the possible effects of membrane voltage fluctuation on response reliability, we performed simulations of leaky integrate and fire neurons with the following parameters: Vthresh: −49mV, Vinit: −70 mV, integration time step = 0.05 ms, Cm = 4.9 ms, gl = .16 us, El = −65. We created a Gaussian shaped tuning curve across 15 different stimuli to define the input current generated by each stimulus. We simulated different levels of membrane voltage fluctuation by adding independent Gaussian noise to membrane voltage at each time step. Noise variance was empirically determined to match values recorded *in vivo* for running and stationary animals. We presented 300ms trials of each stimulus 10 times in a randomized order, and calculated reliability in the same fashion as in the main text.

## Acknowledgements

AJC was supported by a Texas Instruments Stanford Graduate Fellowship; JWP was supported by grants from the McKnight Foundation, Simons Collaboration on the Global Brain (SCGB AWD1004351) and the NSF CAREER Award (IIS-II50186).

## Author Contributions

AJC and JWP conceived the study, AJC performed the analyses, and AJC and JWP wrote the manuscript.

